# Experience alters hippocampal and cortical network communication via a KIBRA-dependent mechanism

**DOI:** 10.1101/2022.05.10.491238

**Authors:** Lilyana D. Quigley, Robert Pendry, Matthew L. Mendoza, Brad. E. Pfeiffer, Lenora J. Volk

## Abstract

Synaptic plasticity is hypothesized to underlie “replay” of salient experience during hippocampal sharp-wave/ripple (SWR)-based ensemble activity and to facilitate systems-level memory consolidation coordinated by SWRs and cortical sleep spindles. However, it remains unclear how molecular changes at synapses contribute to experience-induced modification of network function. The synaptic protein KIBRA regulates plasticity and memory, although its impact on circuit dynamics remains unknown. Here, we recorded *in vivo* neural activity from WT mice and littermates lacking KIBRA to examine circuit function before, during, and after novel experience. In WT mice, experience altered network dynamics in a manner consistent with incorporation of new information content in replay and enhanced hippocampal-cortical communication. However, while baseline SWR features were normal in KIBRA cKO mice, experience-dependent alterations in SWRs were absent. Furthermore, intra-hippocampal and hippocampal-cortical communication during SWRs was disrupted following KIBRA deletion. These results reveal molecular mechanisms that underlie network-level memory formation and consolidation.

## Introduction

Synaptic plasticity plays a critical role in adaptive cognition and memory(*1-6*). In particular, trafficking of AMPA-type ionotropic glutamate receptors (GluAs or AMPARs) within the post-synapse has emerged as a key and conserved regulator of synaptic plasticity and learning(*2, 7*). The synapse-enriched protein KIBRA (<u>Ki</u>dney and <u>Bra</u>in protein, a.k.a. WWC1) is a post-synaptic scaffold that regulates AMPAR trafficking(*8-10*). Accordingly, KIBRA is acutely required in the adult brain for learning and memory as well as synaptic plasticity(*8-11*). Gene variants of KIBRA are associated with normal variation in human episodic and working memory performance(*12, 13*) highlighting its importance in human cognition. In addition, KIBRA and the protein complexes it organizes are implicated in a wide range of neurological/psychiatric disorders known to have synaptic/circuit etiologies(*8, 14*), including autism(*15-17*), schizophrenia(*18, 19*), bipolar disorder(*20*), Tourette’s Syndrome(*21*),and Alzheimer’s disease(*22*). It is unknown how abnormal plasticity may contribute to pathological brain function or how plasticity at individual synapses contributes to larger-scale, circuit-level changes that facilitate memory. Thus, elucidating KIBRA’s role in experience-dependent changes in network function may reveal fundamental links between plasticity, memory, and neurological disorders.

Of particular relevance for the study of network-level memory processes are hippocampal sharp-wave/ripple (SWR) events, brief (50-200 ms), high-frequency (125-300 Hz) oscillations that occur during inactive behavioral states such as slow-wave sleep and quiet wakefulness(*23-25*). SWRs coordinate temporally compressed “replay” of neural activity representing prior or possible future behaviors(*25-28*), and considerable evidence implicates SWRs as a network-level mechanism for memory consolidation and/or retrieval in both rodents and humans(*23, 26, 29-31*). Disruption of SWRs impairs memory(*30, 32-34*) and artificial prolongation of SWRs can enhance memory(*31*), arguing for a causal role of SWRs in mnemonic processes. Further, SWRs are altered in human patients with schizophrenia(*35*) and several animal models of cognitive dysfunction and neuropsychiatric disorders(*36-38*), highlighting their importance in normal brain function.

Of functional relevance to SWR generation by the hippocampal circuit, cortical networks produce brief (0.5-3 s) 10-15 Hz oscillations during slow-wave sleep termed spindles that are strongly associated with cortical plasticity and memory consolidation(*39-42*). The active systems consolidation hypothesis proposes a two-stage model of memory consolidation in which memories are initially encoded in the hippocampus and then transferred to the cortex during sleep(*43*). This process is believed to involve the interaction of SWR events in the hippocampus with sleep spindles in the cortex to support information transfer and memory consolidation(*44, 45*). Indeed, cortical spindles are temporally associated with hippocampal SWRs, and disrupting this relationship impairs sleep-based memory consolidation, arguing that these network-level features represent acute periods of enhanced inter-regional communication required for memory consolidation (*39, 40, 44, 45*), although the synaptic mechanisms which underlie and modulate this coordination are largely unknown.

To study how KIBRA may contribute to circuit-level memory formation, we examined the role of KIBRA in the experience-dependent modulation of hippocampal SWR events and cortical spindles. We evaluated properties of these emergent network features during a novel experience and within rest periods before and after the behavioral session in WT mice and littermates with forebrain-selective deletion of KIBRA from excitatory neurons.

## Results

### Basal hippocampal network dynamics are unaffected by loss of KIBRA

To assess how loss of KIBRA impacts neural circuit mechanisms underlying memory and adaptive cognition, we performed *in vivo* electrophysiological recordings from freely behaving WT mice and littermates lacking KIBRA in excitatory neurons of the forebrain (KIBRA cKO: *Kibra*^*Floxed/Floxed*^ *(8) x* αCaMKII-CreER^T2^ (*46*))(*10*). We simultaneously monitored brain activity in hippocampal areas CA1 and CA3 as well as the anterior cingulate cortex (ACC) before, during, and after exploration of a novel environment with enriched visual and social cues (Fig. 1A, Extended Data Fig. 1). The environment provided a salient experience expected to produce long-lasting changes in synaptic and network function(*3, 47-49*). The total distance traveled, mean velocity, and time spent in active movement were similar between WT and KIBRA cKO mice (Fig. 1B-D), indicating that both groups had equivalent levels of experience.

**Figure 1.**
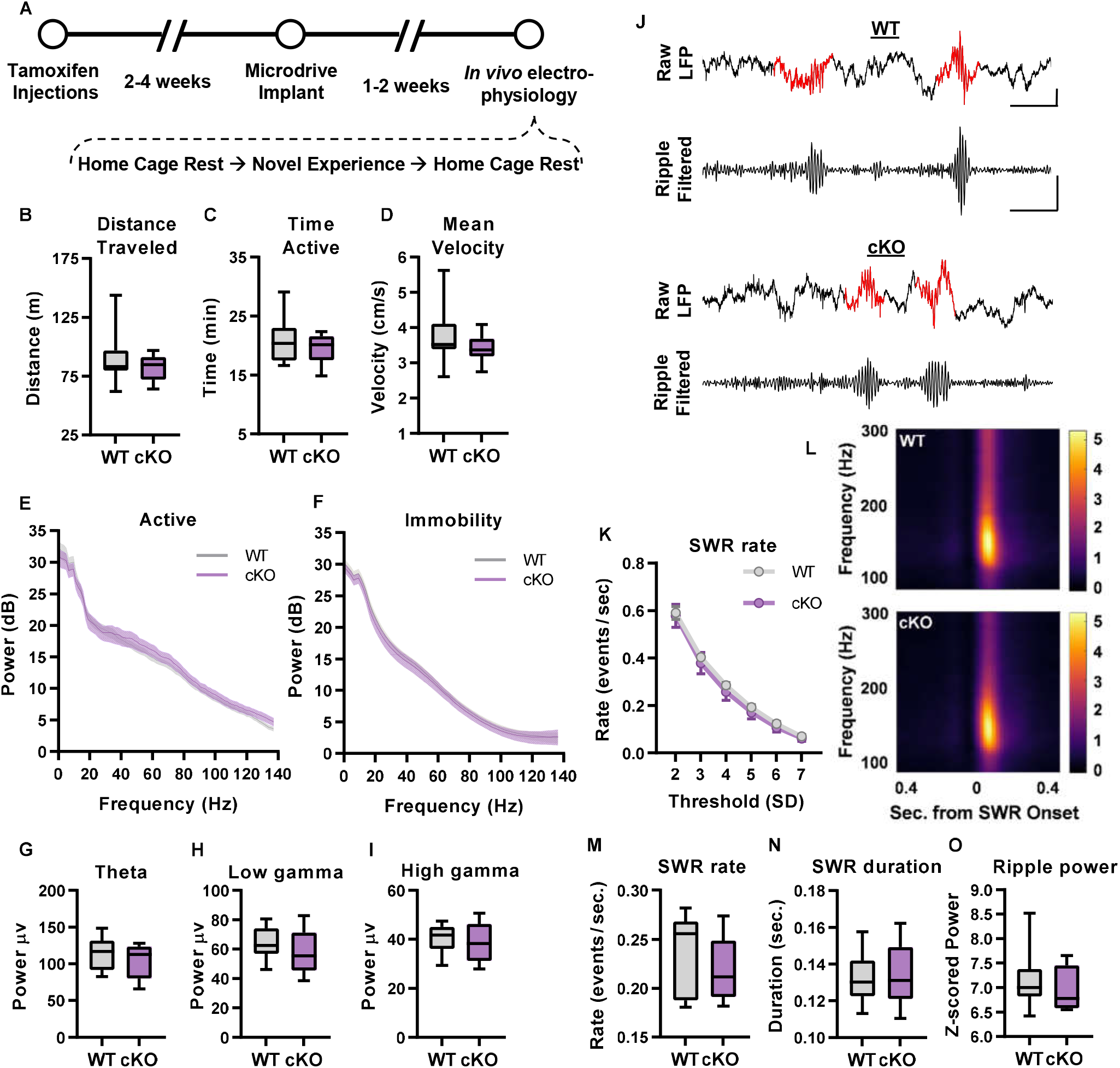
**KIBRA cKO mice show normal baseline hippocampal circuit dynamics prior to novel experience. A, Timeline for experiments. B-D, Total distance travelled (B), total time spent in active movement (C), and mean velocity (D) during behavior sessions. E,F, Baseline power spectral density from pre-experience home cage during active periods (velocity>3cm/s for at least 2.5 seconds, E), and immobility periods (velocity <1cm/s for 3-60s, F). G-I, Mean theta (6-12hz, G) low gamma (25-50Hz, H) and high gamma (65-140Hz, I) band oscillatory power for WT and KIBRA cKO mice during the pre-experience home cage session. J, Example raw and ripple filtered traces depicting detected SWRs in WT (top) and cKO (bottom) mice. Scale bars: 100ms, 200µV, same scale for WT and cKO traces. K, Pre-experience SWR rate by detection threshold (standard deviations above the mean). L, spectrograms depicting increased power in ripple band averaged across all detected SWRs for WT (top) or cKO (bottom) mice, scale bar to right indicates z-scored power. M-O, Baseline ripple properties during pre-experience home cage immobility using detection threshold of 5 SDs above mean SWR power; SWR rate (events/second immobile, M), mean duration of SWR events (N), and peak SWR power (O). Statistics: Unpaired Mann-Whitney Rank sum test (B) or unpaired t-test (C-D, G-I, M-O). Two group test for equal spectrums (E,F). Multiple Mann-Whitney rank sum with Holm-Šidák correction (K). All comparisons were not significant (WT vs cKO, p > 0.1). n=8(cKO)/10(WT) mice per group.**

We first quantified gross aspects of hippocampal network activity by analyzing local field potential (LFP) oscillatory power within area CA1, the primary output of the hippocampus, across several well-studied frequency ranges important for organizing hippocampal information processing and coordinating interregional communication. We observed no differences between genotypes in overall theta, low gamma, or high gamma oscillatory power in the home cage pre-experience (Fig. 1E-I). We then identified SWRs as transient increases in the 125-300 Hz ripple-band power during periods of immobility (Fig. 1J). In the home cage recording prior to novel experience we observed no difference in SWR duration, peak ripple power, or SWR rate at any detection threshold (Fig. 1K-O). Thus, basic network mechanisms of regional synchronization and SWR generation are intact in KIBRA cKO mice, consistent with normal basal synaptic transmission in synapses lacking KIBRA(*8, 10*).

### KIBRA is required for experience-induced changes in SWRs that reflect information updating

Having established that basal circuit properties are grossly normal in KIBRA cKO mice, we next explored whether the loss of KIBRA impacts experience-dependent changes in circuit-level hippocampal function. The occurrence rate of SWRs has been shown to transiently increase following salient experience(*50-52*), an effect that is blocked by systemic pharmacological inhibition of NMDA receptors(*53*). Given KIBRA’s critical role in synaptic plasticity(*8-10*), we hypothesized that changes in SWR drive following experience would be reduced in KIBRA cKO mice. However, both WT and KIBRA cKO mice displayed similar experience-dependent modulation of SWR rate (Fig. 2A, B).

**Figure 2.**
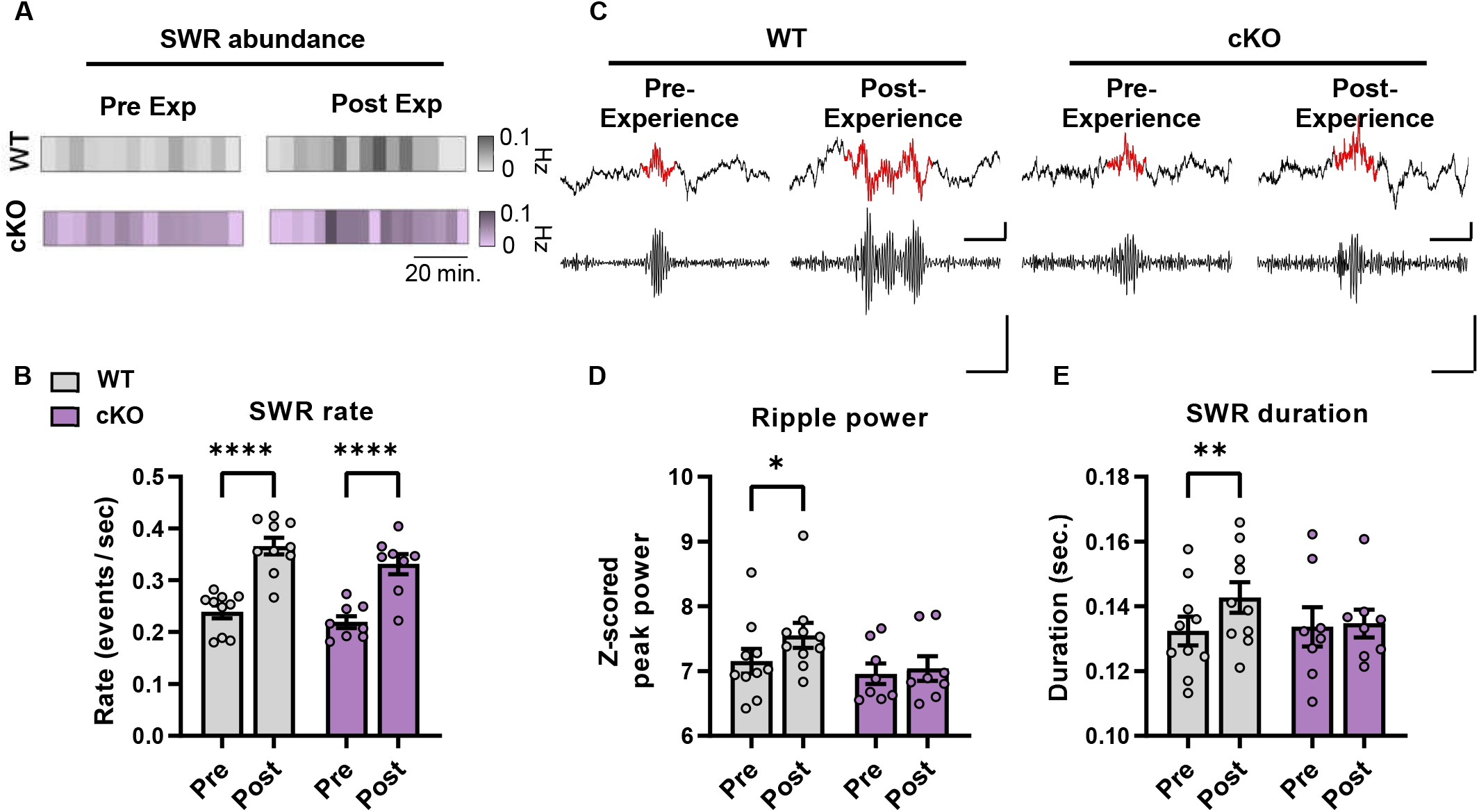
**Experience-dependent modulation of CA1 SWR properties is impaired in KIBRA cKO mice, while modulation of ripple abundance is intact. A, Pre- and post-experience SWR abundance shown in 5 min segments for a representative WT (grey, top) and cKO (purple, bottom) mouse. B, SWR rate (events/sec during immobility) in pre- and post-experience home cage sessions (main effect of experience p < 0.0001, pre vs. post WT p < 0.0001, pre vs. post cKO p < 0.0001). C, Example trace showing experience dependent increases in SWR power and duration in WT but not cKO mice, Scale bar 100ms, 200mV. D, Peak SWR power during pre- and post-experience home cage sessions (main effect of experience p = 0.0284, pre vs. post WT p = 0.0168, pre vs post cKO p = 0.8335). E, SWR duration during pre- and post-experience home cage sessions (main effect of experience p = 0.0279, experience x genotype p = .0656, pre vs. post WT p = 0.0091, pre vs. post cKO p = 0.9476). Statistics: Two-way RM ANOVA with Šidák’s multiple comparisons. ****p<0.0001, ***p<0.001, **p<0.01, * p<0.05. n=8(cKO)/10(WT) mice per group.**

Notably, this increase in pre- to post-experience SWR drive was independent of the threshold used to detect SWRs (Extended Data Fig. 2). These findings suggest that KIBRA expression in excitatory hippocampal neurons is unnecessary for experience-dependent changes in SWR drive and argue that the source for this plastic change resides elsewhere.

We next assessed whether KIBRA expression was necessary for experience-dependent changes in the information content of SWRs. Experience dramatically changes the representation of information within SWR-based replay(*54-56*). Like the experience-dependent change in SWR drive, incorporation of novel information into SWRs can be blocked by systemic pharmacological inhibition of NMDAR-dependent synaptic plasticity(*56*). Neuronal synchrony, which contributes to oscillatory power, also increases during SWRs after salient experience(*24, 50, 57*). In addition, replays representing longer spatial trajectories are encoded by SWRs of longer duration(*58*), and long-duration SWRs are increased during memory tasks(*31*). Thus, we hypothesized that if features of experience were incorporated into SWR-based replay via a KIBRA-dependent plasticity mechanism, we would observe an increase in the duration of SWRs and the power of the ripple component of SWRs in WT mice but not in mice lacking KIBRA expression. Supporting our prediction and consistent with prior data in rats, WT mice displayed a significant increase in both ripple power and SWR duration in the post-experience rest period (Fig. 2C-E). However, ripple power and SWR duration were unchanged following novel experience in KIBRA cKO mice (Fig. 2C-E), consistent with impairments in synaptic plasticity following KIBRA loss(*8-10*).

During SWRs, hippocampal networks are coordinated via the 25-50 Hz low gamma oscillation(*55, 59*). Low gamma power is transiently increased in hippocampal area CA1 during SWRs and the representation of information during SWR-based replay is temporally segmented by hippocampal low gamma(*55, 59*). We observed a transient increase in low gamma power during SWRs of both WT and KIBRA cKO mice (Fig. 3A-E). During pre-experience home cage periods, there was no significant difference in SWR-based gamma power between genotypes (Fig. 3F), in agreement with a lack of basal changes in synaptic function and SWR properties in KIBRA cKO mice(*8, 10*) (Fig. 1J-O).

**Figure 3.**
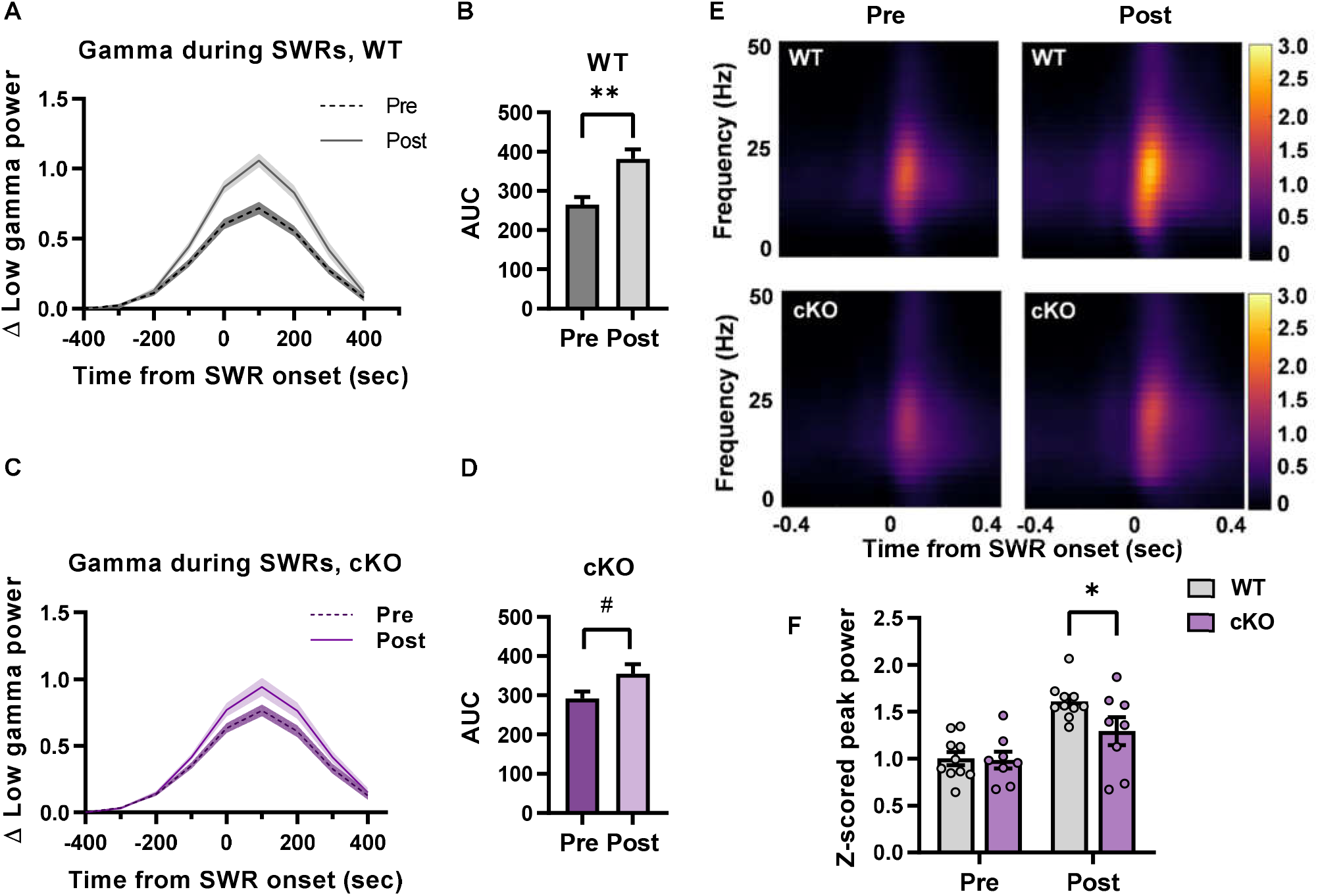
**Experience-dependent increase in SWR-associated low gamma power is impaired in KIBRA cKO mice. A C, Transient increase in low gamma power during SWR events in pre- and post-experience home cage for A, WT and C, cKO mice. B,D, Area under curve (+/- 450ms surrounding SWR onset) for SWR-associated low gamma power in B, WT (p=0.0019) and D, cKO (p=0.0528) mice. E, Time-frequency spectrograms showing experience modulation of SWR-associated low gamma power in WT (top) and cKO (bottom) mice, scale bar to right indicates z-scored power. F, Peak low gamma power during SWRs (main effect of experience p <0.0001, experience x genotype interaction, p = 0.0095, pre WT vs. cKO p = 0.9916, post WT vs.cKO p = 0.0469). Statistics: Two-way RM ANOVA with Šidák’s Multiple comparisons (F), unpaired t-test (B,D). **p<0.01, * p<0.05, # 0.05<p<0.1. n=8(cKO)/10(WT) mice per group.**

When we compared pre-experience SWRs to post-experience SWRs, WT mice displayed an experience-dependent increase in low gamma power, consistent with increased network synchrony and incorporation of novel information into SWR-based replay (Fig. 3A, B, E, F). However, KIBRA cKO show impaired experience-dependent modulation of low gamma power during SWRs (Fig. 3C-F), providing further support for the hypothesis that plasticity deficits following KIBRA loss impair incorporation of salient information content during SWR-based replay.

SWRs that occur during quiet wake are both functionally and mechanistically distinct from those that arise during slow-wave sleep(*33, 60*). The above analyses combined all SWRs during home cage recording sessions irrespective of behavioral state (quiet wakefulness vs. sleep). To determine whether KIBRA deletion selectively impacts a distinct subpopulation of SWRs or instead produces a more general impairment in SWR modulation, we defined epochs of putative sleep vs. wakeful rest based on immobility duration and hippocampal oscillatory dynamics(*61*). We observe no difference in overall sleep durations, rest durations or gross sleep architecture across genotypes (Extended Data Fig. 3). WT mice display a similar experience-dependent increase in SWR duration, power, and rate for SWRs occurring in both sleep and quiet wakefulness (Fig. 4), indicating that the mechanisms underlying these plastic changes in circuit dynamics are observed across behavioral states and likely reflect broad changes in hippocampal network function. The duration and power of SWRs in KIBRA cKO mice do not change from pre- to post-experience rest, regardless of sleep or wake state (Fig. 4A-D). Interestingly, while the rate of SWRs significantly increased across experience in both sleep and wake for cKO mice, the experience-dependent increase in SWR rate was significantly attenuated in the quiet wake state, such that the overall ripple rate in post-experience wake was reduced in KIBRA cKO mice compared to WT littermates (Fig. 4E). In general, however, these data demonstrate that KIBRA loss has similar effects on SWRs occurring in both sleep and quiet wakefulness.

**Figure 4.**
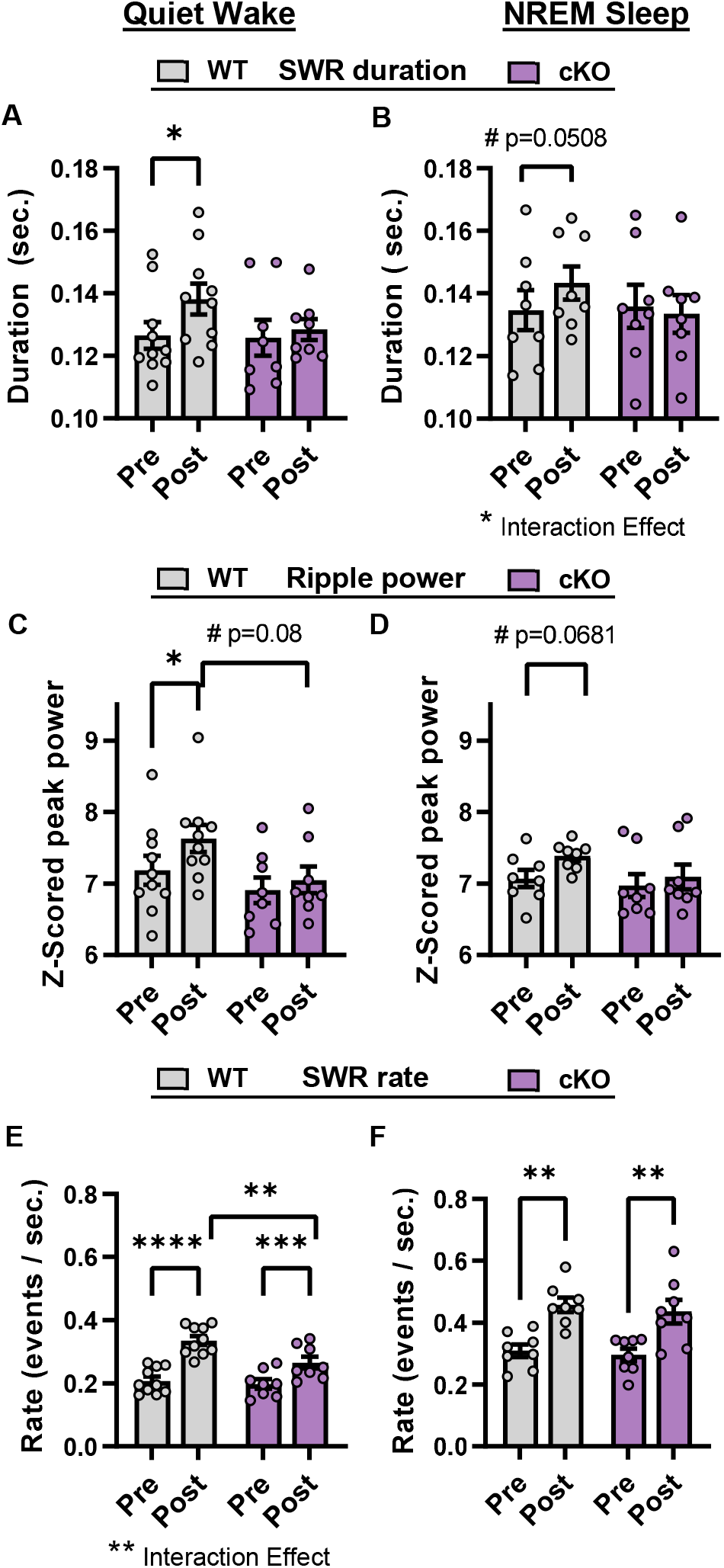
**Loss of KIBRA impairs experience-modulation of remote SWR properties during quiet wake and sleep. A, C, E, SWR properties for SWRs detected in quiet wake during pre- and post-experience home cage sessions, including A, duration (main effect of experience p=0.0176, WT pre vs. post p=0.0104, cKO pre vs. post p=0.7695) C, peak power (main effect of experience p=0.0182, WT pre vs. post p=0.0177,cKO pre vs. post p=0.6401), and E, rate (main effect of experience p<0.0001, experience x genotype p=0.007, main effect of genotype p=0.0507, WT pre vs. post p<0.0001,cKO pre vs. post p=0.0006, WT vs. cKO pre p=0.8775, post p = 0.0048), B,D,F, properties of SWRs detected during putative non-REM sleep (NREM sleep), including B, duration (experience x genotype p=0.0464, WT pre vs. post p=0.0580, cKO pre vs. post p=0.7666, WT vs. cKO pre p=0.9877, post p =0.4660), D, peak power (main effect of experience p=0.0376, WT pre vs. post p=0.0681,cKO pre vs. post p=0.6140, WT vs. cKO pre p=0.8466, post p=0.2629), and F, rate (main effect of experience p<0.0001, WT pre vs.post p=0.0010, cKO pre vs. post p=0.0017, WT vs. cKO pre p=0.9214, post p =0.8073. Statistics: Two-way RM ANOVA with Šidák’s multiple comparisons (A-F). ****p<0.0001, ***p<0.001, **p<0.01,* p<0.05, # 0.05<p<0.1. A,C,E, n=8 (cKO), n=10 (WT) mice. B,D,F, n=8 (cKO), n=8 (WT) mice.**

### Deficits in SWRs begin to emerge during experience

The above results indicate that novel experience generates plastic changes in the hippocampal network that are reflected in altered SWR features during post-experience rest, and that KIBRA is necessary to either induce or maintain those changes. To dissociate these two possibilities, we examined SWRs which occurred during brief pauses within the novel experience itself (“on-task” SWRs). Because incorporation of task-relevant information into on-task replay occurs rapidly(*25*) and requires synaptic plasticity(*56*), we hypothesized that WT mice should display modification of SWRs during behavior, whereas such modifications should be impaired or absent in KIBRA cKO mice. Indeed, when we quantified features of SWRs arising during the novel experience, we observed that KIBRA cKO mice had significantly weaker ripple power than WT littermates (Fig. 5A). In addition, KIBRA cKO mice displayed a trend toward decreased SWR rate (Fig. 5B). These data are consistent with a role for KIBRA in rapid, plastic changes across the hippocampal network during experience, changes which normally persist into post-experience periods of memory consolidation. Unexpectedly, SWR duration and SWR modulation of low gamma power were similar across genotypes for on-task SWRs (Fig. 5C-E), with differences in these measures emerging in post-experience rest (Figs. 2-4).

**Figure 5.**
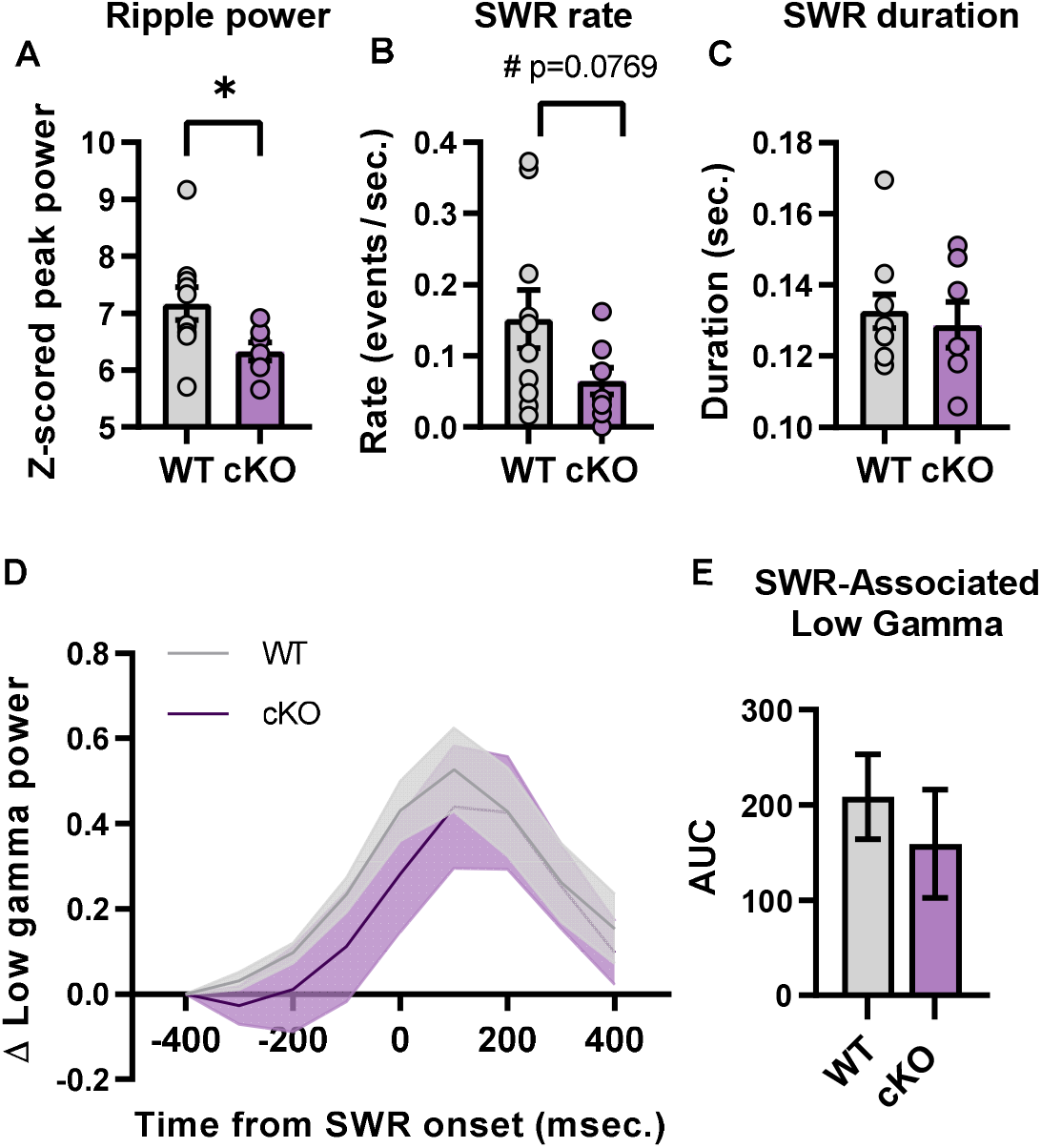
**Loss of KIBRA impairs awake SWR properties during novel experience. A, Z-scored peak ripple power (p= 0.0377), B, SWR rate (p=0.0769), and C SWR duration (p=0.6240), for SWRs with ripple power greater than 3 S.D. above the mean that occurred in periods of immobility during a novel experience. D, SWR-associated increase in low gamma power. E, Area under the curve for SWR associated low gamma power (+/- 450ms surrounding SWR onset), (p=0.4968). Error bars reflect mean ± SEM. Statistics: Unpaired t-test (A-C,E) with Welch’s correction (C). * p<0.05, # 0.05<p<0.1 n=7-8 (cKO, one mouse had no on-task SWRs and was excluded from A,C-E), n=10 (WT) mice.**

### Intrahippocampal coordination during SWRs requires KIBRA

During SWR-based replay, hippocampal areas CA3 and CA1 are tightly synchronized to facilitate information transfer(*59*). CA3 input to CA1 is coordinated by the low gamma oscillation(*62*) and is crucial for organized, sequential activation of CA1 place cells during SWRs(*63*). Given prior work demonstrating a role for KIBRA in plasticity at CA3→CA1 synapses(*8-10*) and our data demonstrating that KIBRA cKO mice show impaired experience-dependent modulation of SWR-associated low gamma power (Fig. 3), we hypothesized that coordination between CA1 and CA3 would be disrupted after loss of KIBRA. In both WT and KIBRA cKO mice, ripple power and low gamma power were transiently elevated in area CA3 during CA1-identified SWRs in pre-experience rest (Extended Fig. 4), and no difference was observed between genotypes in the basal CA3 low gamma power outside of SWRs (Fig. 6A). Similar to area CA1, area CA3 of WT mice displayed an experience-dependent increase in both ripple power and SWR-associated low gamma power, an effect that was abolished in KIBRA cKO mice (Fig. 6B,C), indicating that KIBRA loss produced similar impairments in experience-dependent changes in both CA1 and CA3 networks.

**Figure 6.**
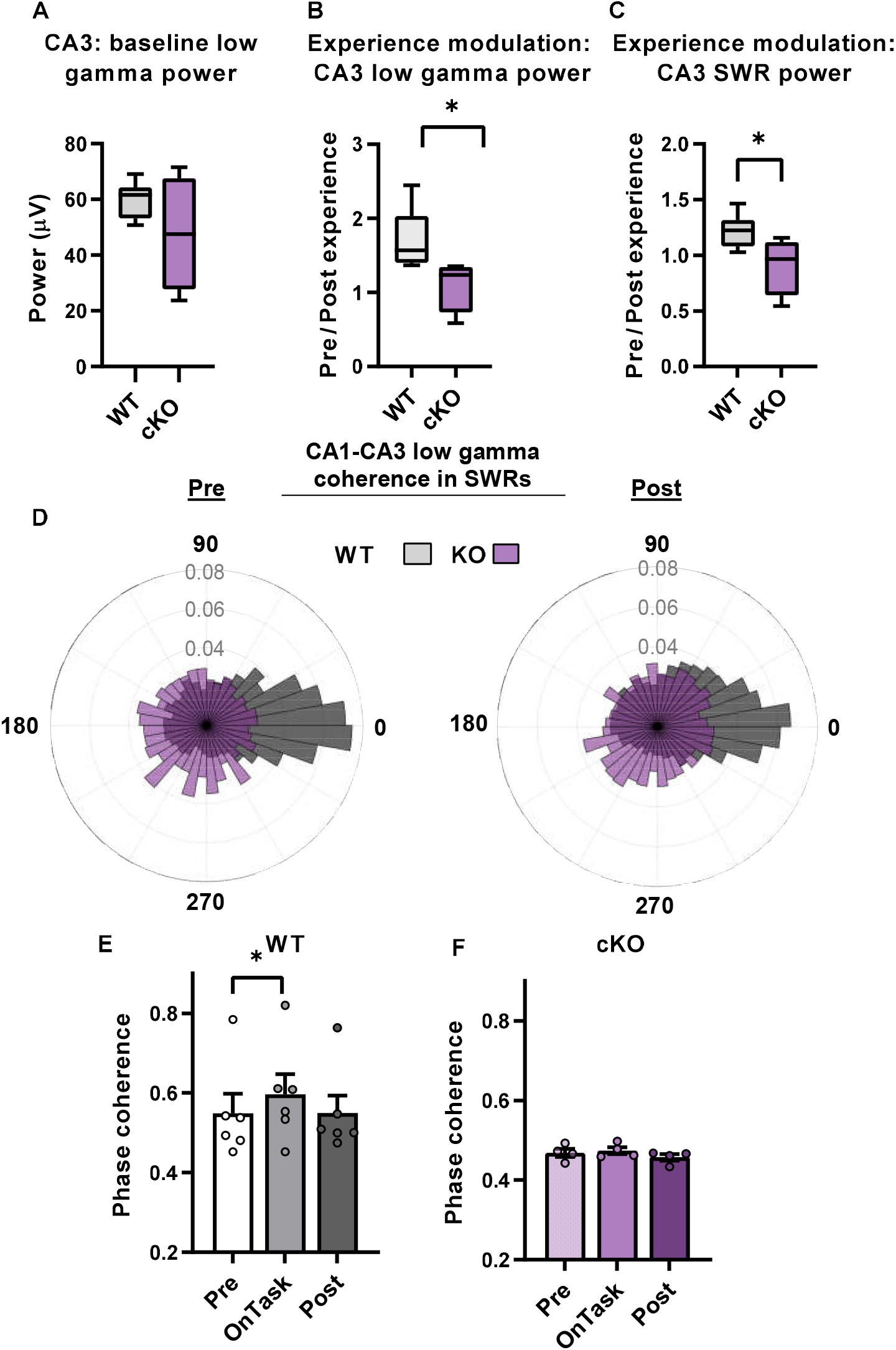
**Loss of KIBRA alters experience modulated CA3-CA1 communication during SWRS. A, Baseline CA3 low gamma power in pre-experience rest (p=0.3185). B, Experience modulation of peak CA3 low gamma power during identified CA1 SWRs, shown as ratio of post/pre-experience (p=0.0411). C, Experience modulation of peak CA3 ripple power during CA1 SWRs shown as ratio of post-/pre-experience (p=0.0428). D, Normalized polar histogram (36 bins, 10*°* each) showing distribution of CA3-CA1 SWR-associated low gamma phase offsets during pre- (left) and post-experience (right) home cage sessions for all WT (gray) and KO (purple) SWRs. Bars represent probability (Rayleigh Test for uniformity: WT pre z=306.76 p <0.0001, post z=546.33 p <0.0001; cKO pre z=18.19 p < 0.0001, post z=1.3542 p=0.2582. ANOVA, main effect of experience p < 0.0001 and genotype p <0.0001, experience x genotype interaction p<0.0001, Kuiper’s Two-Sample Test: WT vs. KO distribution Pre p=0.003, Post p=0.002). E,F, Mean CA3-CA1 phase coherence (ISPC) during SWRs by session for WT (E), and cKO (F). Statistics: Unpaired t-test (A-C), with Welch’s correction (A). Rayleigh test for uniformity, Harrison-Kanji Circular ANOVA, Kuiper Two Sample Test with Bonferroni-Holm corrections for multiple comparisons (D). Friedman test with Dunn’s correction for multiple comparisons (E,F). WT n=6, cKO n=4 mice (A-C, E-F). n = # ripples in D, WT: Pre (n=3894), Post (n=7432), cKO: Pre (n=2262) Post (n=3002). ****p<0.0001, **p<0.01, * p<0.05.**

We next examined the precise temporal synchronization between CA1 and CA3 during SWRs. As low gamma oscillations are known to facilitate communication between CA3 and CA1(*62*) and discretize information during SWR-based replay(*55*), we examined phase-phase coupling of the low gamma oscillation across CA3 and CA1 networks. We observe striking low gamma coherence between CA1 and CA3 in WT mice during SWRs before and after experience (Fig. 6D). However, KIBRA cKO mice display severely disrupted synchronization, both in pre- and post-experience SWRs (Fig. 6D). Additionally, during on-task SWRs we find a significant increase in low gamma synchrony in WT mice that is not present in KIBRA cKO mice (Fig 6E,F, Extended Data Fig. 5). Together these findings demonstrate that temporal coordination of CA3-CA1 activity at the low gamma frequency is disrupted during SWRs in the cKO at basal levels and fails to undergo experience-dependent changes.

### Experiences synchronizes hippocampal SWRs and cortical spindles during subsequent NREM sleep and requires intact KIBRA expression

To assess the role of KIBRA in systems-level memory consolidation, we examined whether the coordination of ACC sleep spindles with CA1 SWRs is altered by the loss of KIBRA. We find no significant differences between WT and cKO mice in the sigma band power used for spindle detection (Fig. 7A-B) or basal spindle properties in pre-experience rest, including spindle rate, duration, or power (Fig. 7C-E). We find that spindle density and duration increase with experience in WT mice, consistent with prior findings(*64, 65*) (Fig. 7F,G). However, while KIBRA cKO mice exhibit an increase in spindle rate, they fail to show experience-dependent changes in cortical spindle duration (Fig. 7F,G). Interestingly, while we did not observe experience-dependent increases in power across all ACC spindles in WT or cKO mice (Fig. 7H), when we restricted our analysis to ACC spindles occurring during identified hippocampal SWRs, we observed a significant experience-induced increase in spindle power in WT mice that was not present in cKO mice (Fig. 8A). Several studies have demonstrated coupling of hippocampal SWR and neocortical spindle events during post-learning sleep(*39, 44, 66*). Given the impairments in cross-regional (CA3-CA1) synchronization observed in KIBRA cKO mice (Fig. 6), we predicted that hippocampal-cortical coordination might also be altered after loss of KIBRA. Therefore we examined time lags from SWRs to the nearest spindle and found a shift closer to zero lag from pre- to post-experience in WT mice (Fig. 8B,D), indicating enhanced synchronization of these two brain regions. However, this improvement in inter-regional coordination is absent in KIBRA cKO mice (Fig. 8C,D). As SWRs tend to be nested within the troughs of spindles(*66*), we next examined the properties of SWRs that were nested inside spindles. The proportion and peak ripple power of SWRs nested within spindles increased after experience in WT but not cKO mice (Fig. 8E,F). Finally, we found that the number of cortical spindles which overlapped with multiple consecutive hippocampal SWRs increased across experience in WT but not cKO mice (Fig. 8G). Importantly, this is not a trivial result of increased SWR and spindle rate, as both WT and KIBRA cKO mice show elevated SWR (Fig, 2A,B) and spindle (Fig. 7F) rate. Together these findings indicate that KIBRA is necessary for experience-dependent enhancement of SWR-spindle coordination.

**Figure 7.**
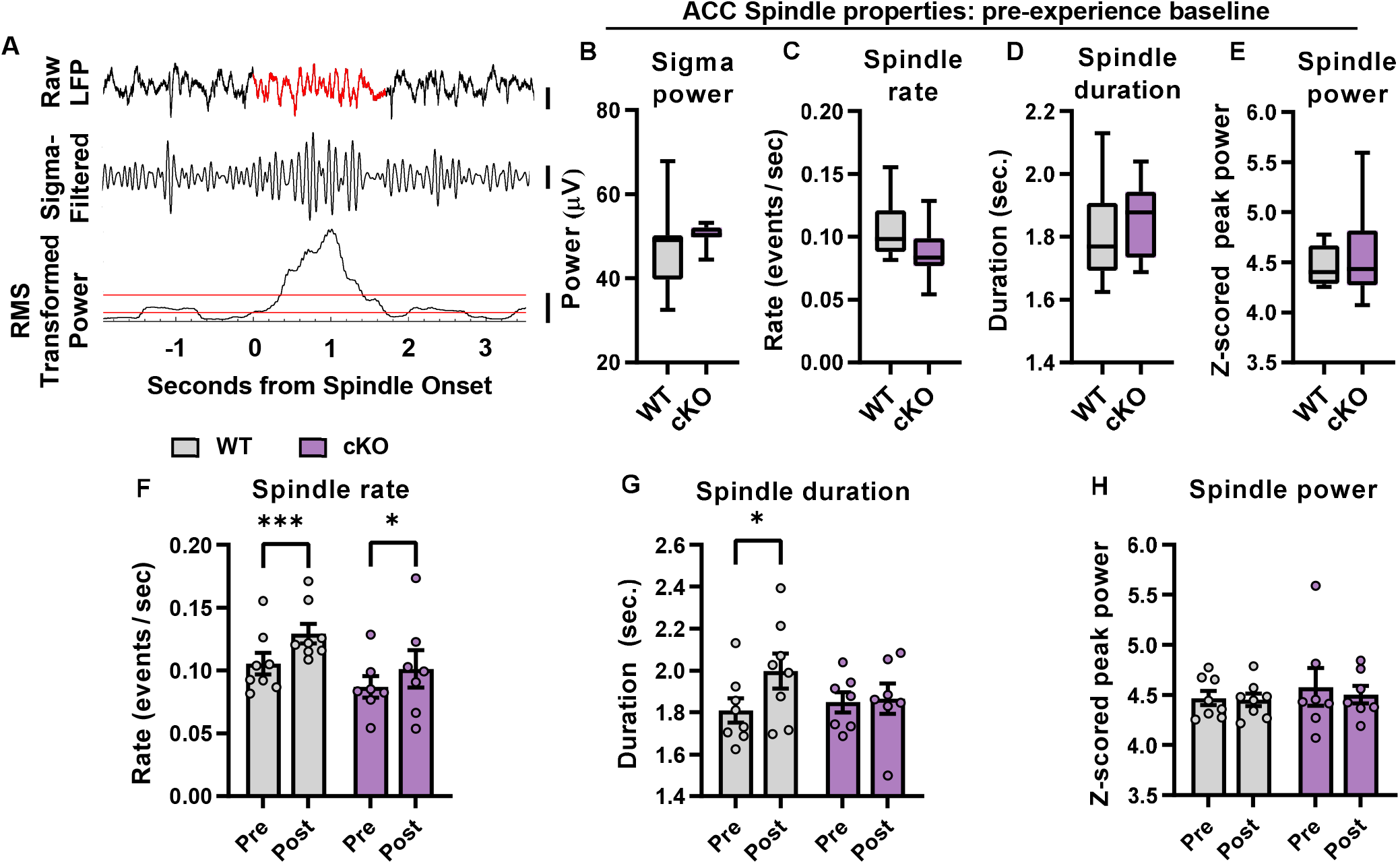
**Impaired experience-dependent regulation of ACC spindles in KIBRA cKO mice. A, Example of a spindle event detected in the ACC. Top: raw LFP during spindle event, middle: sigma-filtered trace (10-15Hz), bottom: RMS transformed trace used for spindle detection. Scale bars: raw LFP (500µV), sigma-filtered (200µV), transformed (1×10^6^ µV^3^). B, ACC sigma power in pre-experience home cage sessions during all immobility periods (p = 0.1416). C-E, Pre-experience baseline properties of identified spindles; C, rate (# of spindle events per sec) (WT vs. cKO, p=0.1571), D, mean spindle duration (WT vs. cKO, p=0.6135), E, peak sigma power during spindles (WT vs. cKO, p=0.8865). F-H, Experience modulation of ACC spindles; F, spindle rate (main effect of experience p=0.0002, WT pre vs. post p =0.0009, cKO pre vs. post p=0.0453), G, spindle duration (main effect of experience p=0.0631, WT pre vs. post p=0.0343, cKO pre vs. post p=0.9676), H, peak sigma power across all spindles (WT pre vs. post p=0.9729, cKO pre vs. post p=0.6787). Statistics: Unpaired t-test (C,D). Mann Whitney Rank Sum (B,E). Two way RM ANOVA with Šidák’s multiple comparisons (F-H). n=8-9(WT, one mouse had no detected post-experience sleep and was excluded from F-H)/n=7(KO) mice per group. ***p<0.001,* p<0.05.**

**Figure 8.**
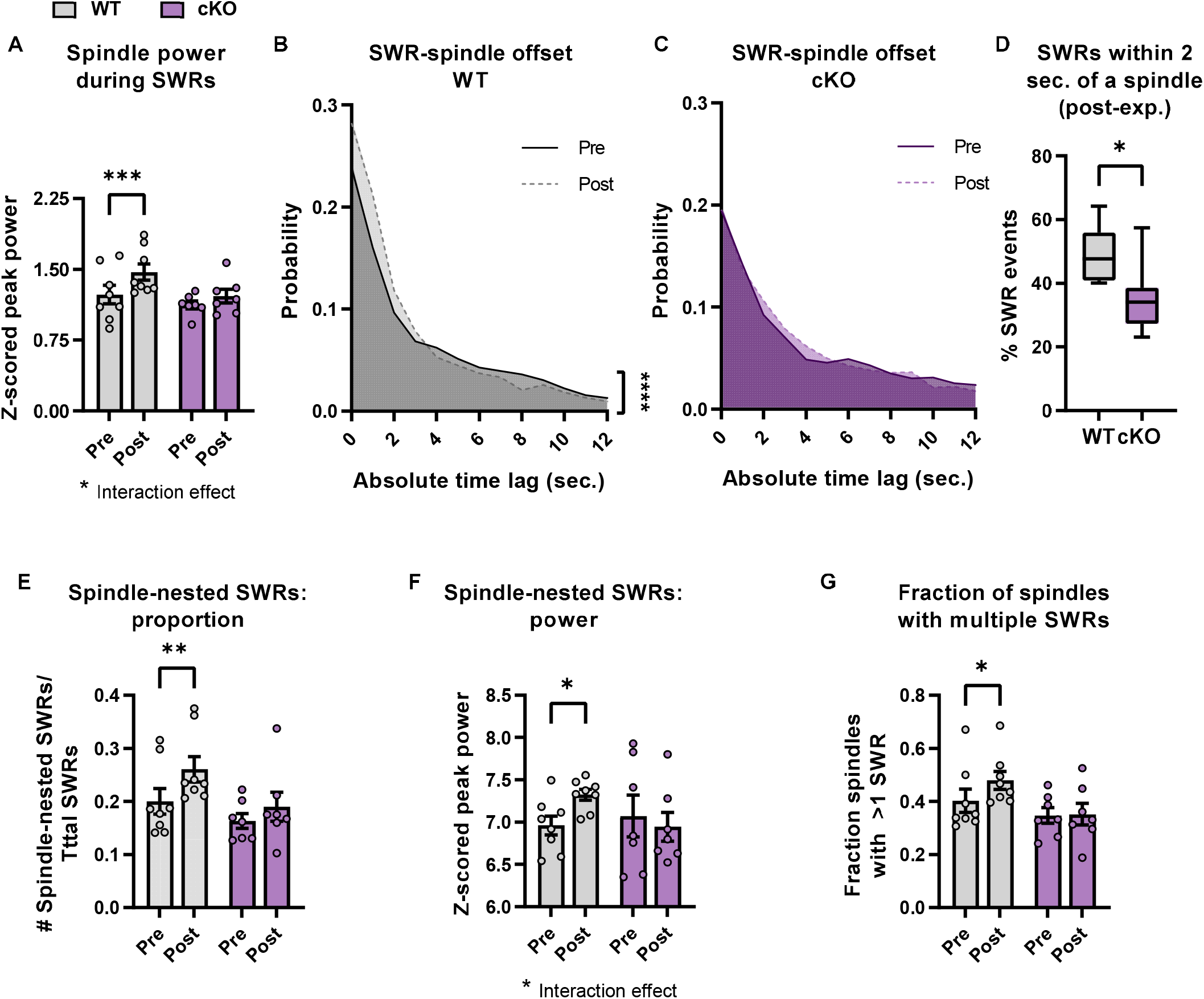
**Coordination of ACC spindles and CA1 SWRs during sleep is disrupted by loss of KIBRA. A, Peak ACC sigma power for spindles co-occurring with CA1 SWRs (main effect of experience p=0.0002, experience x genotype p=0.0376, WT pre vs. post p=0.0002, cKO pre vs. post p=0.1250). B, C, Probability distribution for absolute lag between CA1 SWRs and ACC spindle onset during putative NREM sleep for WT(B) and cKO (C), pre vs. post: WT p < 0.0001, cKO p= 0.1063. D, Percent of SWR events occurring within 2 seconds of a spindle during post-experience rest (WT vs. KO, p=0.0217). E, Proportion of ACC spindles that co-occur with CA1 ripples (main effect of experience p=0.0007, WT pre vs. post p=0.0013, cKO pre vs. post p=0.1694). F, Peak ripple power of SWRs nested within spindles (experience x genotype p = 0.0308, WT pre vs. post p=0.0408, cKO pre vs. post p = 0.6521). G, Fraction of spindles containing more than one nested SWR (# spindles containing >1 SWR / # spindles containing 1 or more SWRs), (main effect of experience p=0.0425, experience x genotype p=0.0684, main effect of genotype p=0.0964, WT pre vs. post p=0.0168, cKO pre vs. post p=0.9806). Statistics: Two way RM ANOVA with Šidák’s multiple comparisons (A, E-G). Wilcoxon Rank-Sum test on distributions with Bonferroni-Holm correction for multiple comparisons(B,C). Unpaired t-test (D) n=8(WT)/n=7(KO) mice per group (A, D-G). n = # ripples in B,C, WT: pre(n=3969) post(n=5262) cKO: pre(n=3352) post (n=4555). ****p<0.0001,***p<0.001,**p<0.01,* p<0.05.**

## Discussion

KIBRA plays a key role in regulating AMPAR trafficking, synaptic plasticity, learning and memory(*8-11*). Substantiating this crucial role are numerous human genetic studies that implicate KIBRA in memory and disorders of complex cognition(*12, 13, 16, 18, 21, 67*). However, whether and how KIBRA-dependent cellular mechanisms impact *in vivo* network dynamics was previously unknown. Here, we examined SWRs before, during, and after salient novel experience in adult mice lacking the synapse-enriched protein KIBRA. Considerable evidence implicates SWR-based replay as a memory consolidation and retrieval mechanism in both rodents and humans. Thus, SWRs serve as a quantifiable measure of circuit-level memory formation and consolidation(*23, 24, 26, 29, 30, 33*). We find that under pre-experience baseline conditions SWRs are grossly normal following removal of KIBRA, consistent with our previous findings that basal synaptic transmission is unaffected in these mice(*10*). However, loss of KIBRA abolishes experience-dependent changes in several SWR properties, including duration, ripple-band power, and SWR-associated gamma-band power. In addition, KIBRA cKO mice display impaired basal intra-hippocampal (CA3-CA1) coordination at the low gamma frequency during SWRs and fail to produce experience-dependent changes in intra-hippocampal and hippocampal-cortical communication during SWRs. These experiments were performed in a conditional knockout mouse line in which KIBRA was selectively deleted from mature, predominantly excitatory forebrain neurons(*10, 46*), arguing against altered neuronal development as an explanation for these observations. Thus, these results collectively indicate that KIBRA is acutely required for experience-dependent SWR modulation and that KIBRA-dependent mechanisms facilitate synchronization across interregional networks. Furthermore, while the current study was not designed to provide behavioral assessment of long-term spatial memory, these data further argue that KIBRA-dependent changes in sequential replay and hippocampal-cortical communication during SWRs may contribute to memory impairments resulting from loss of KIBRA(*8-11, 22*).

Novel experience has been shown to increase SWR drive in post-experience rest periods(*50, 51, 53, 68*). *In vitro* slice experiments demonstrate that stimuli which establish LTP can facilitate the spontaneous generation of SWRs(*69*). Consistent with a role for synaptic plasticity in experience-dependent changes in SWR expression, the increase in SWR drive is abolished by pharmacological blockade of NMDARs during the experience(*53*). The experience-dependent increase in SWR drive observed in both WT and KIBRA cKO mice in our study indicates that the site of NMDAR-dependent plasticity which underlies this increased drive is unlikely to reside in synapses onto excitatory neurons of the hippocampus. Rather, this plasticity may occur at CA3 inputs onto CA1 inhibitory neurons. Indeed, both experimental and computational work suggest that excitatory input from CA3 pyramidal neurons onto local parvalbumin-positive (PV^+^) basket cells may trigger SWR initiation via generation of high-frequency oscillatory firing of inhibitory populations(*70, 71*). Our findings therefore argue that excitatory input onto PV^+^ inhibitory neurons is a likely site of experience-dependent increases in SWR drive.

In addition to changes in SWR drive, considerable evidence supports a role for synaptic plasticity across the hippocampal network in the incorporation of experience-relevant information in SWR-based replay. The activity patterns of hippocampal pyramidal cells with overlapping firing fields produce NMDAR-dependent LTP between those cell pairs(*49*), thus increasing the probability for those neurons to co-fire in subsequent offline SWRs(*24, 57*). In agreement with these findings, pharmacological inhibition of NMDAR function during experience prevents coherent representation of the current experience in subsequent SWR-based replay, indicating that NMDAR function is important for establishing synaptic plasticity during experience which is later revealed in the experience-specific content of SWR-based replay (*56*). Changes in the information content of SWRs following novel experience are reflected in increased overall firing rate and ripple power(*51, 57*). In addition, SWRs that replay longer virtual trajectories have a longer duration(*58*), and long-duration SWRs are increased during memory tasks(*31*).

Thus, experience-dependent changes in SWR duration and ripple power are likely both driven by synaptic plasticity. However, NMDAR blockade broadly affects multiple forms of synaptic plasticity across many cell types. While we did not record from the large single unit populations necessary to unambiguously quantify “replay” content, we observe increased ripple power and SWR duration following salient experience in WT mice whereas KIBRA cKO mice fail to show such experience modulation of SWRs, supporting the hypothesis that incorporation of recent experience into hippocampal replay requires KIBRA-dependent synaptic plasticity mechanisms within the excitatory pyramidal neurons of the hippocampal network.

The information content in SWR-based replay is temporally organized by the low gamma oscillation(*55*), the power of which transiently increases during SWRs(*38, 55, 59*). Gamma activity during SWRs is strongly impaired in an age-dependent fashion in a mouse model of Alzheimer’s disease(*38*), suggesting a role for SWR-embedded low gamma in mnemonic processes. While stronger gamma coherence across the CA3-CA1 networks during SWRs is predictive of the presence of coherently structured replay(*59*), it was previously unknown whether experience directly impacted the low gamma-SWR relationship. Here, we show for the first time that the transient increase in gamma activity during SWRs is significantly elevated following a novel experience. Hippocampal neurons are phase-locked to low gamma during SWRs(*55, 59*) and experience increases overall firing rates during SWRs(*57*); thus, the enhancement in gamma power we observe during SWRs in WT mice is consistent with these prior findings. Although KIBRA cKO mice display a normal low gamma/SWR relationship during pre-experience rest periods, the absence of KIBRA impairs experience-dependent changes in this relationship, suggesting that recruitment of gamma phase-locked firing in SWRs following novel experience may be impaired in the absence of KIBRA.

While synaptic plasticity during experience can influence subsequent SWRs, the firing patterns within SWRs can themselves produce LTP(*3*), which is believed to facilitate information transfer from hippocampal to cortical networks and underlie long-term, systems-level memory consolidation(*43*). Supporting this model, blocking SWRs during post-experience rest impairs long-term memory(*30, 32, 72, 73*). During SWRs, the hippocampal and cortical networks become more tightly synchronized(*44, 74*). In particular, cortical sleep spindles are temporally coupled to hippocampal SWRs(*39, 40, 44, 45*). Disrupting this relationship impairs sleep-based memory consolidation, while enhancing SWR-spindle coupling improves memory(*39, 40, 44, 45*). We observe that the duration of cortical spindles and the strength of SWR-associated cortical spindles increases across experience in WT mice, but not in KIBRA cKO mice. Furthermore, we observe an experience-dependent enhancement of the temporal coupling between hippocampal SWRs and cortical spindles in WT, but not cKO mice. While the present study was not designed to address prolonged, multi-day plastic changes across the hippocampal or cortical networks, we hypothesize that the lack of experience-dependent changes in SWR-spindle coordination likely results in impaired memory consolidation and may account for the long-term memory impairments observed in mice lacking KIBRA expression(*8, 10, 11*).

In contrast to SWR properties and SWR-spindle coupling, which were not different between WT and KIBRA cKO mice in pre-experience conditions, CA3-CA1 communication during SWRs was abnormal in cKO mice even during this baseline period. Consistent with prior work in rats(*59*), we observe strong coherence across hippocampal areas CA3 and CA1 on a low gamma timescale during SWRs in WT mice. This coherence was significantly weaker in cKO mice, demonstrating impaired communication between these regions. Given the lack of basal changes in synaptic function observed in KIBRA cKO(*8, 10*), it is unclear what accounts for this impaired communication. Both areas CA1 and CA3 display similar overall gamma power and a normal SWR-based increase in gamma power is observed prior to novel experience in KIBRA cKO mice; thus, the lack of low gamma coherence is unlikely to be a result of impairments in the mechanisms of gamma generation. Rather, the lack of coherence may result from a shift in the balance of CA1 drive from CA3 inputs relative to layer 3 entorhinal cortex (EC3) inputs(*61, 62, 75*).

It remains unknown how experience modifies the hippocampal network to allow for specific sequences to be re-expressed in SWR-based replay or how specific plasticity mechanisms regulate the expression of these sequences during different stages of memory acquisition and consolidation. Our data are consistent with a model in which novel or salient experience drives rapid KIBRA-dependent synaptic plasticity across the hippocampal and cortical networks, and that these plastic changes fundamentally alter the properties and content of SWR-based hippocampal replay and facilitate enhanced hippocampal-cortical communication during subsequent sleep. As KIBRA is known to regulate AMPAR trafficking and expression at excitatory synapses(*8-10*), the lack of experience-dependent changes in SWR function and inter-regional coordination are likely a result of impaired synaptic plasticity at these sites, however our data cannot rule out contributions from other neuronal functions of KIBRA. Our findings are consistent with the hypothesis that synaptic plasticity at excitatory CA3 synapses onto PV^+^ basket cells may facilitate experience-dependent increases in SWR drive, while plasticity at excitatory synapses on CA1 pyramidal neurons is likely responsible for changes in SWR content. As KIBRA, plasticity, and SWRs are commonly linked to cognitive and neuropsychiatric disorders(*14, 18, 21, 36-38, 67*), additional work investigating how these mechanisms converge to regulate information processing will provide critical mechanistic insights into both normal and pathological brain function.

## Materials and Methods

### Experimental Design

The primary objective of this study was to investigate if and how disrupting a specific memory-related synaptic plasticity mechanism (KIBRA-mediated AMPAR regulation) affects the ability of hippocampal and cortical networks to adapt to novel experience. To accomplish this goal, we recorded neural activity from hippocampal areas CA1 and CA3 and anterior cingulate cortex of freely behaving mice lacking KIBRA expression in CaMKIIα+ (excitatory) forebrain neurons and wild type littermates before, during and after a novel experience.

### Subjects

The University of Texas Southwestern Institutional Animal Care and Use Committees approved all animal protocols in this study. Male mice ages 2.5-5 months were used for all experiments. A tamoxifen-inducible deletion strategy was used to selectively remove KIBRA from mature, excitatory forebrain neurons. KIBRA cKO (CaMKIIα-CreERT2*+*:*KIBRA*^flox/flox^) and WT littermate controls (CaMKIIα-CreERT2***-***:*KIBRA*^flox/flox^) mice of C57Bl/6N genetic backgrounds were generated by crossing floxed *KIBRA* mice (*KIBRA*^*flox/flox*^) with hemizygous CaMKIIα-CreERT2 transgenic mice, which express CreERT2 (Cre recombinase fused to a mutated estrogen receptor) under the *CamkII*α promoter(*46*). KIBRA cKO and WT mice between 2.5 and 4 months of age were injected with tamoxifen (IP, 100mg/kg, 2x/day, 5 days), resulting in ∼70% reduction of KIBRA protein in the hippocampus of KIBRA cKO mice(*10*). Mice were allowed to recover for at least two weeks after the last tamoxifen injection before microdrive implantation.

### In vivo recordings

Mice were surgically implanted with custom-made tetrode microdrives (4-4.5g fully assembled) using previously reported surgical procedures(*55*). Microdrive components were designed using Solidworks software, printed using a 3D printer (Formlabs), and assembled using methods adapted from(*26*). Each microdrive array contained 8 independently adjustable tetrodes (twisted bundles of four 17.8 µm 90% platinum/10% iridium wires, California Fine Wire) bilaterally targeting (in mm) ACC (ML: ±.4-.5 AP: +.5 DV: 1.25), CA1 (ML: ±1.5 AP: -2.28 DV: 1.5), CA3 (ML: ±2.56 AP: -2.08 DV: 2.25) and a reference in overlying cortex. The tetrode tips were electroplated with gold solution (Neuralynx) to reduce the impedance to 150-200 kΩ before surgery. Small jewelry screws (Component Supply) and dental cement were used to anchor microdrives to the skull of the animal. One anchor screw behind lambda, overlying the cerebellum, served as a ground. Subjects were implanted under regulated isoflurane-oxygen anesthesia and fixed in a flat-skull position in a stereotaxic frame. For a majority of recordings, hippocampal tetrodes were individually lowered to their final recording locations over 1-2 weeks until ripples and hippocampal spike activity were identified. Hippocampal tetrodes for a subset of mice (3 WT, 2 cKO) were lowered to the target recording depths during surgery. No differences were observed between these groups in either recording quality or experimental results. ACC tetrodes were lowered to the target recording depths during surgery. Subcutaneous Buprenorphine (0.006 mg/ml at 0.01 ml/g body weight) was given as a postoperative analgesic. Post-surgery, all implanted animals were individually housed on a 12 hr light/dark cycle with ad libitum access to chow and water in their home cages.

At the end of experiments, mice were given a lethal dose of Euthasol, and electrolytic lesions were made to confirm recording locations. Mice were then transcardially perfused with ice-cold sterile saline followed by 4% paraformaldehyde. Postmortem electrode locations were verified by examining 75µm brain coronal sections made with a Precisionary Instruments Compresstome (VF-300).

### Data acquisition

Data were collected using a Neuralynx (Bozeman, MT) data acquisition system. Recording electrodes were connected to a Digital Lynx recording system via an HS-36 headstage and tether. During electrophysiological recordings the animal’s behavior was monitored by overhead video recorded continuously at 30 FPS and synchronized to electrophysiological data acquisition (Cheetah software). The animal’s position was determined using LEDs or infrared tracking stickers positioned at the front and back of the microdrive array (pointing towards nose or tail, respectively). Analog neural signals were digitized at 32,556 Hz. Continuous local field potential (sampled at 3,255.6 Hz) were recorded using Cheetah data acquisition software (Neuralynx).

### Behavior

Mice were handled for at least 5 min on each of 5 consecutive days prior to experiments. Recordings were conducted at least 7 days post-implantation during the animal’s light cycle to ensure adequate sleep in the pre- and post-experience rest periods. Continuous electrophysiological recordings were conducted throughout the following: a 1-hour pre-experience home cage session in which subjects were allowed to freely behave in the home cage, mice were then transferred to the center chamber of a novel 40 cm x 60 cm plexiglass box (22 cm tall) with four interior baffles (22 cm tall, angled 10-14.5 cm long) that protruded from long-axis walls and provided visual and tactile cues as mice explored the space. High contrast visual cues were also present on the walls in the room where the recording was performed. Following a 20-minute exploration of the novel arena, a novel con-specific male mouse and a novel object were placed into the chamber (10 minutes) after which the novel object was replaced by an additional novel con-specific mouse (10 minutes). The unimplanted mice providing social salience were kept in a small wire mesh cage (10 cm tall, 9 cm diameter) which prevented them from causing damage to the recording apparatus on the implanted subject, but allowed for close interaction. The novel behavioral experience was followed immediately by a 1-hour post-experience home cage session. Animal positions were quantified from behavior videos via *deeplabcut(76)*.Velocity was used to define behavior states, including active movement (velocity >3cm/s), immobility (<1cm/s), and sleep (<1 cm/s for at least 60s).

### LFP Analysis

Analyses were performed using custom MATLAB scripts following accepted analysis methodology. Reference signals were digitally subtracted from hippocampal electrodes prior to analysis. For each tetrode, one representative electrode from each region was selected for local field potential analyses. Theta (6-12 Hz), low gamma (25-50 Hz), high gamma (65-140 Hz), and ripple (125-300 Hz) bands were isolated from the raw LFP by bandpass filtering in the respective frequency bands. Power was defined as the absolute value of the smoothed (Gaussian kernel, sigma = theta 300 ms; low gamma 85ms; high gamma 30ms; ripple 12.5 ms) Hilbert transform of this filtered signal. 0°/360° was defined as the trough of the filtered oscillation. Removal of 60 Hz line noise was done using spectrum interpolation. Power spectral densities were generated via Chronux using the multitaper method.

SWR-triggered spectrograms were computed using the short-time Fourier transform with 200ms temporal bins, 190 ms overlap, and 2hz frequency resolution. A z-score was computed for each frequency band using the mean and standard deviation of the LFP power during the pre-experience home cage session from CA1. Spectral power was averaged for all SWRs for each animal.

### SWR associated gamma power and coherence

SWR associated low gamma power was quantified using 100ms time bins. The change in low gamma power during SWRs was quantified by subtracting the baseline (−450 to -350 preceding each SWR) power from each time point. Gamma synchrony was computed during SWRs by determining the phase offset between CA1 and CA3 tetrodes for all time points during a SWR and using the inter-site phase clustering method (ISPC)(*77*). ISPC is defined as the length of the average of phase angle difference vectors from two electrodes. An ISPC value was calculated for each SWR and averaged per animal. For polar histograms, the mean phase offset was calculated for each SWR and values were combined to obtain a distribution of gamma phase offsets for all SWRs. Change in gamma coherence from baseline was measured by calculating the ISPC for phase offset distributions in each 100 ms bin, ±450ms from the onset of each SWR and subtracting the baseline value.

### Ripple Detection and analysis

Sharp-wave/ripples were detected on CA1-targeted tetrodes as peaks above the mean power (5 SD above the mean unless otherwise noted), in the smoothed (Gaussian, sigma 12.5 ms) ripple band (125-300 Hz) power, during only periods of immobility (velocity < 1 cm/s). The start and end of each ripple were defined as the point when the smoothed ripple power crossed the mean+1SD. For all SWR events, thresholds were set using the pre-experience home cage session to calculate the mean and SD. SWR rate was calculated as the number of detected events divided by the number of seconds spent immobile. Power and duration were calculated for each SWR event and then values were averaged for all events per subject.

### NREM and quiet wake definition

Slow-wave sleep was defined as times of prolonged immobility (>60 seconds) when the ratio of theta-to-delta power was below a threshold set by a blinded observer based on visual inspection of the CA1 spectrograms (0-20Hz), using a 2 second window with 0.2 second advancement. To calculate the theta-to-delta ratio, a moving average of the envelope power was taken for theta (5s window) and delta (12.5s window). After calculating the ratio, the data was z-scored. For analyses of both pre and post experience sleep, only mice with detected sleep for both sessions were used for analysis.

### Spindle detection and analysis

Spindle events were detected in the ACC during putative NREM sleep periods as described(*78*) using increases in the processed, cubed root-mean-square (RMS) transformed sigma power (10-15Hz). Detection thresholds for each mouse were set using the RMS transformed sigma power during quiescent periods in the pre-experience home cage session. Briefly, an upper threshold was used to identify peaks in the cubed RMS signal and a lower threshold was used to define the start and ends. Spindle events shorter than 0.5 s or longer than 10 s were eliminated from analysis. Events with an inter-event interval less than 0.05 s were combined. Spindle rate was calculated as the number of detected events divided by the time spent in NREM sleep. Power and duration were calculated for each spindle event, and then values were averaged for all events per subject. Temporal coordination of ACC spindles with CA1 SWRs was estimated by finding the offset between each NREM SWR and the nearest spindle start time. Probability distributions were generated for each genotype using the absolute value of each ripple-spindle time lag.

### Statistical Analysis

Unless otherwise noted, data from each mouse were averaged such that each mouse contributed one data point for each comparison. All data were assessed for normality using the Shapiro-wilk or Lilliefors test, in addition to visual inspection of distributions and QQ plots. Equality of variance between groups was assessed using either an F test or Spearman’s test, and examination of residual homoscedasticity plots. One or two-way ANOVAs with post hoc multiple comparison tests or t-tests were used to evaluate significance. Statistical tests were chosen based on sample size, hypotheses, and checks of statistical assumptions. In the small number of instances when assumptions were not met for the above tests, either nonparametric statistics were used, or the data was normalized by calculating a ratio of post/pre-experience values. All data are shown as mean ± SEM for bar graphs, or median + min-to-max for box and whisker plots unless otherwise indicated. Statistical analyses were carried out with GraphPad Prism software or custom analysis code in MATLAB. For Power Spectral Densities, a two-group test for equal population spectrums was conducted via Chronux(*79*). The Circular Statistics toolbox was used to analyze phase angle data.

## Author contributions

Conceptualization: LJV, LQ

Methodology: LQ, RP, LJV, BEP

Resources: LQ, RP, MLM

Investigation: LQ, RP

Formal Analysis: LQ, BEP

Visualization: LJV, LQ

Supervision: LJV, BEP

Writing—original draft: LJV, LQ

Writing—review & editing: LJV, BEP, LQ

## Data and materials availability

All data, code, and materials will be made available upon request to the corresponding author.

