## Supplemental Data for "Experience alters hippocampal and cortical network communication via a KIBRA-dependent mechanism"

**This file includes:**

Figs. S1 to S5

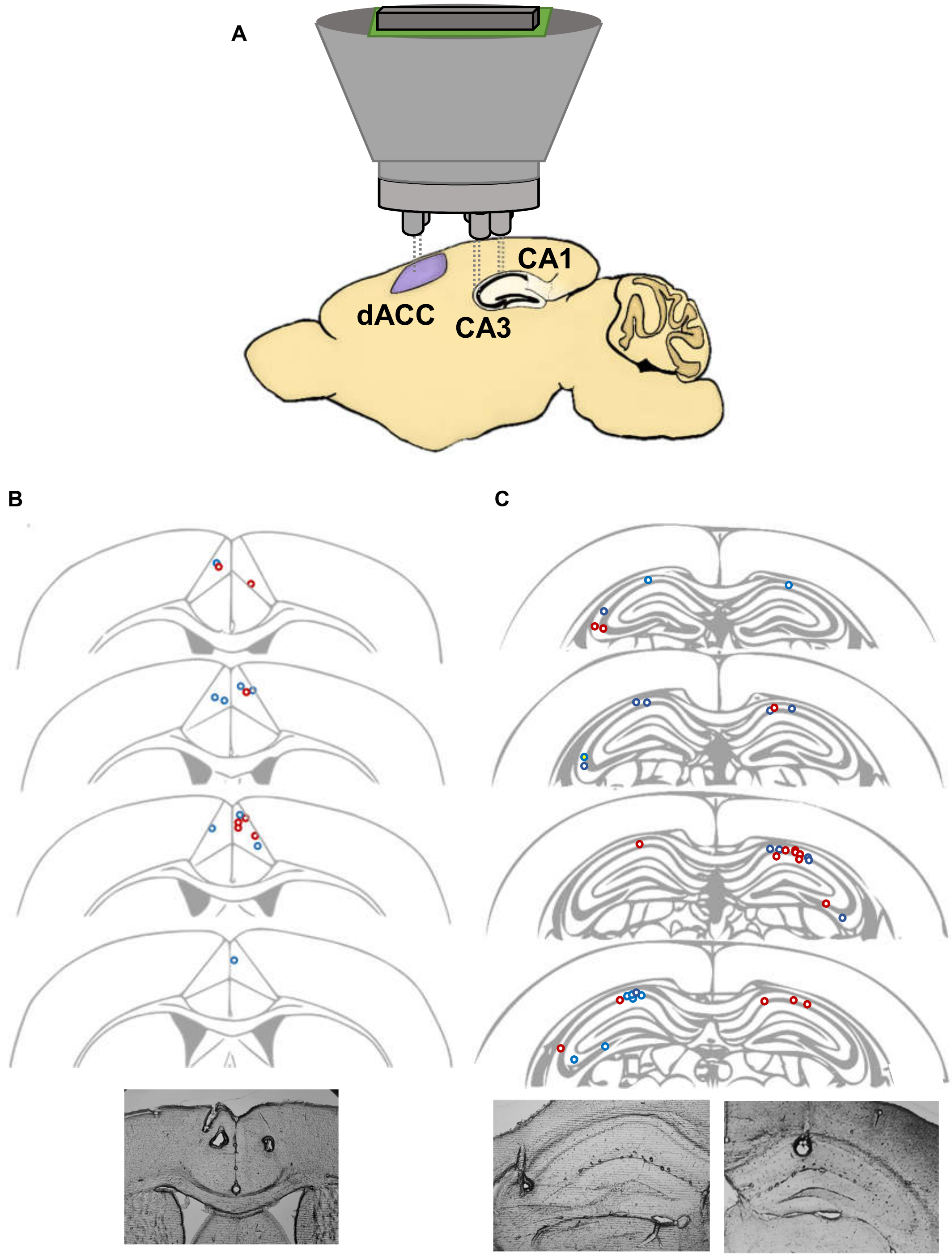


**Figure S1. A**, Schematic showing brain regions bilaterally targeted by tetrode microdrive. **B**, Top: map of ACC recording locations used for analysis shown by genotype as blue open circles (WT) and red open circles (cKO), bottom: image of electrolytic lesions in ACC from one animal. **C**, Top: map of hippocampal recording locations used for analysis from CA1 and CA3, shown by genotype as blue open circles (WT) and red open circles (cKO), bottom: electrolytic lesions from CA1(left) and CA3(right).

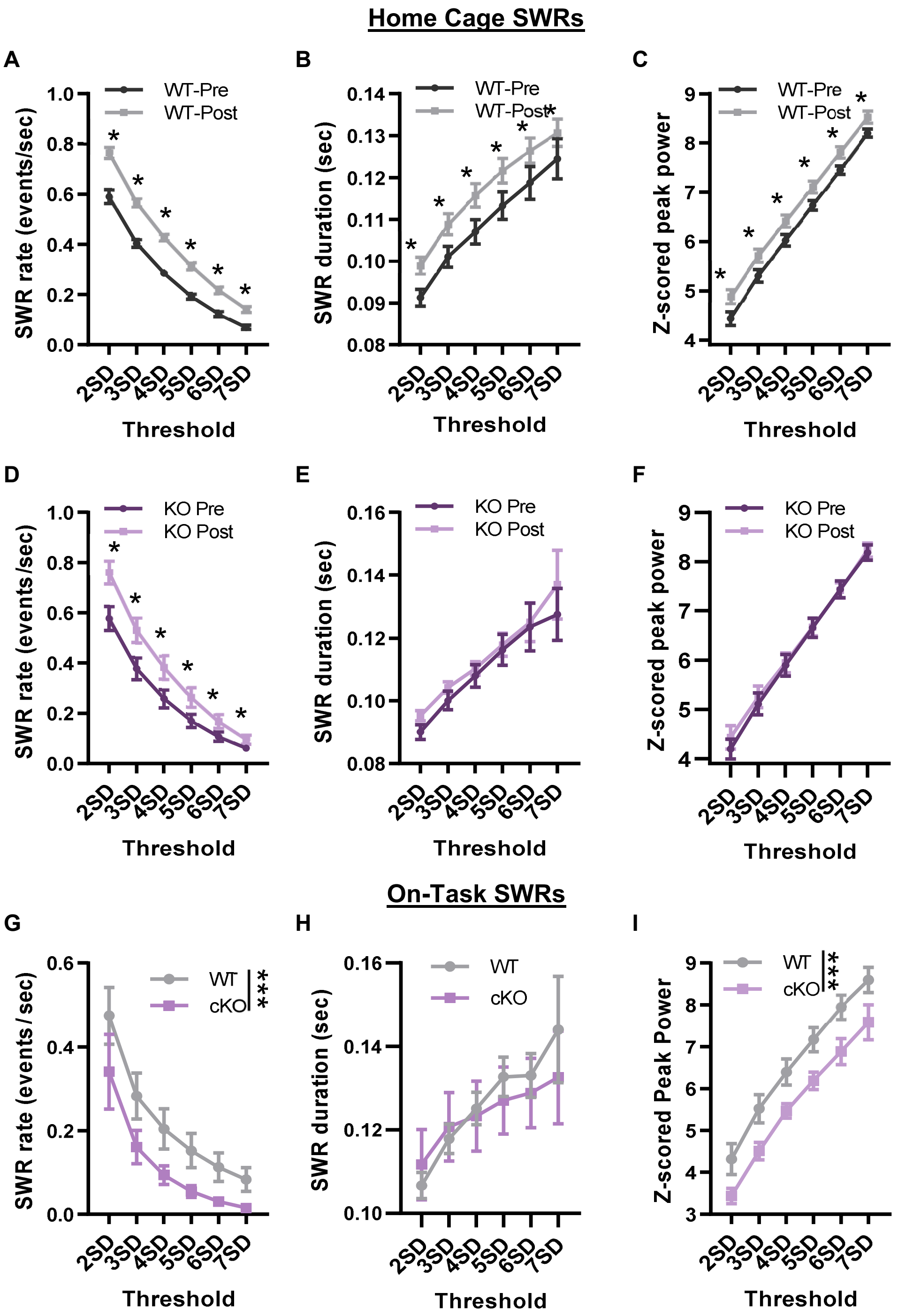


**Figure S2. SWR properties by detection threshold**. **A-F,** Ripple rate (**A,D),** duration (**B,E**), and peak power (**C,F**) by threshold (standard deviations above the mean, SD) during home cage (pre and post experience home cage periods for WT (**A-C**) and KIBRA cKO (**D-F**) mice. **G-I**, properties for SWRs occurring during novel experience (On Task) for WT and cKO mice. **G**, SWR rate: main effect of genotype p = 0.0002, **H**, duration: main effect of genotype, p = 0.5597, **I**, power: main effect of genotype p <0.0001. Statistics: Wilcoxon matched-pairs Signed rank test with Holm-Šidák correction (A-F) Ordinary Two-way ANOVA(G-I). n=8(cKO) n=10(WT) mice

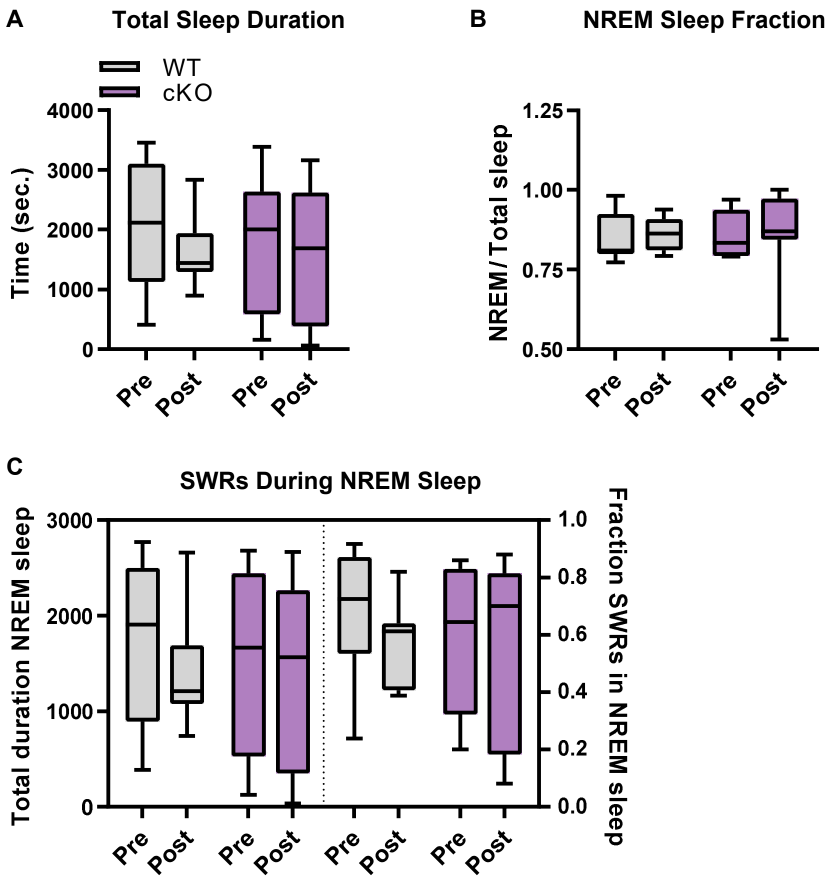


**Figure S3. Sleep properties and proportion of home cage SWRs by state.** **A**, Total duration (seconds) of putative sleep detected in pre- and post-experience home cage sessions. **B,** Proportion NREM sleep relative to total sleep time. **C**, Total duration of detected NREM sleep (left panel) and fraction of SWRs that occurred in NREM sleep vs. NREM sleep + quiet wake (right panel). WT vs. cKO not different significantly different ($p > 0.1$) for any comparison. Statistics: Two-way RM ANOVA with Šidák's multiple comparisons (A-C). n=8(cKO) n=8 (WT) mice.

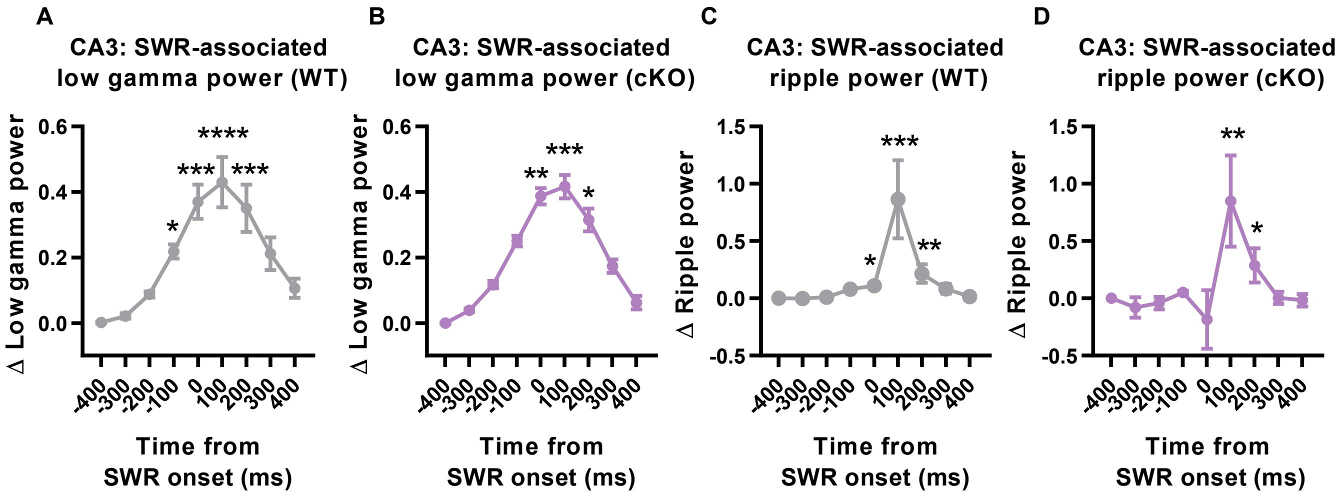


**Figure S4. CA3 Ripple and low gamma power during CA1 SWRs . A-B,** Transient increase in CA3 low gamma power during CA1-identified SWR events in pre-experience, shown as change from baseline (-450ms) for **A** (WT), and **B** (cKO), **C-D,** Transient increase in CA3 ripple power during CA1-identified SWR events in pre-experience for **C,** WT and **D,** cKO**.** Statistics: Friedman test with Dunn's multiple comparisons ****p<0.0001,***p<0.001,**p<0.01,* p<0.05. n=4 (cKO), n=6 (WT) mice.

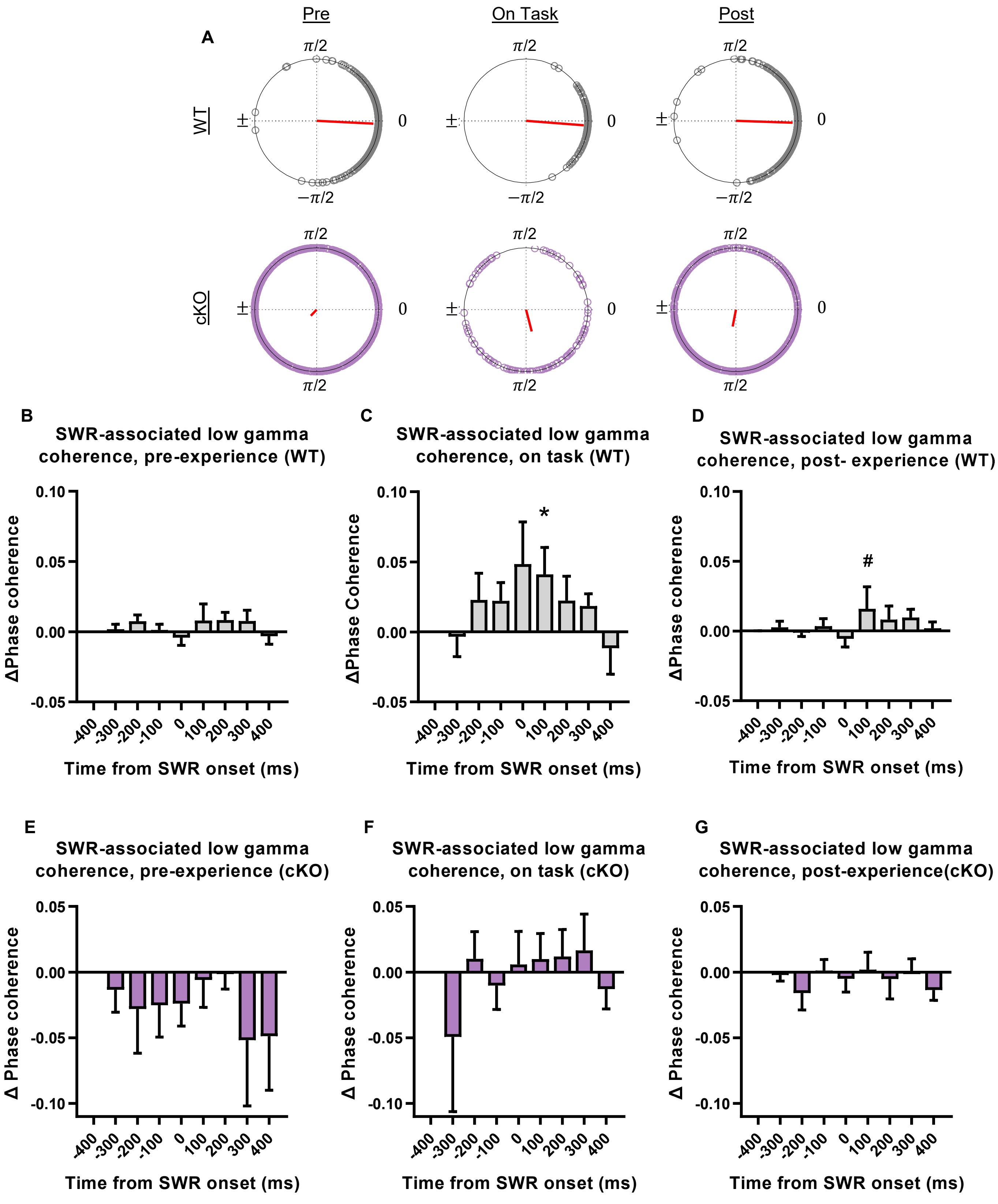


**Figure S5. CA1-CA3 communication. A,** Example circular dot plot for one representative WT (top, grey circles) and KIBRA cKO (bottom, purple circles) mouse showing CA1-CA3 low gamma phase offsets during each CA1-identified SWR (each dot represents mean offset during one SWR). Red line depicts mean resultant vector length. **B-C**, Change in CA1-CA3 low gamma synchrony from baseline (-450ms from ripple onset) for pre-experience (**B,E**) , on-task (**C,F**), and post-experience. Statistics: Friedman test with Dunn's multiple comparisons (**B-G**), * $p<0.05$, # $0.05<p<0.1$. n=4 (cKO), n=6 (WT) mice.
